# 4-Methylpyrazole-mediated inhibition of Cytochrome P450 2E1 protects renal epithelial cells, but not bladder cancer cells, from cisplatin toxicity

**DOI:** 10.1101/2024.11.10.622845

**Authors:** Jephte Y. Akakpo, Erika Abbott, Benjamin L. Woolbright, Anup Ramachandran, Rick G. Schnellmann, Darren P. Wallace, John A. Taylor

## Abstract

Cisplatin is an effective chemotherapeutic drug for the treatment of bladder cancer, though cisplatin-induced nephrotoxicity (CIN) occurs in approximately 20-30% of patients, limiting its clinical use. Evidence has shown that cytochrome P450 2E1 (CYP2E1), a drug metabolism enzyme expressed in proximal tubules, mediates the production of reactive oxygen species (ROS) during cisplatin-induced injury. Previously, we showed that the repurposed drug 4-methylpyrazole (4MP; fomepizole) blocks CYP2E1 activity and prevents acetaminophen-induced liver injury. Here, we investigated the potential protective effects of 4MP against CIN. Male and female C57BL/6J mice were treated with a single 20 mg/kg dose of cisplatin for 3 days (acute) or 9 mg/kg/week for 4 weeks (repeated dosing regimen) with or without 50 mg/kg 4MP as a co-treatment. Our findings revealed that acute treatment with cisplatin induced severe histological tubular damage and elevated plasma BUN and creatinine levels in male mice, but not in female mice. This difference correlated with higher basal CYP2E1 expression in the kidneys of male mice compared to female mice. We also found that cisplatin increased renal CYP2E1 activity and that inhibition of CYP2E1 with 4MP significantly reduced cisplatin induced cell death in male mice and primary normal human kidney cells. By contrast, human bladder cancer cells do not express CYP2E1, and treatment with 4MP did not interfere with cisplatin’s anti-cancer effects in human bladder cancer HTB9 cells. This study highlights the critical role of CYP2E1 in CIN and suggests that its inhibition with 4MP in the kidney is a potential prophylactic therapeutic option to prevent CIN in bladder cancer patients without affecting its anti-neoplastic effect.

**Impact Statement:** This study demonstrates that cytochrome P450 2E1 (CYP2E1) is a critical mechanistic target in the prevention of cisplatin-induced nephrotoxicity (CIN). It also indicates that CYP2E1 plays an important role in mediating sex-specific differences in CIN. Finally, this study reveals that targeting CYP2E1 with 4methylpyrazole offers a promising prophylactic approach to reducing CIN in clinical settings while preserving the anti-cancer efficacy of cisplatin.

## INTRODUCTION

Cisplatin is the single most actively used drug in the management of solid tumors, including Bladder cancer (BCa) (Chestnut et al., 2021; Kamat et al., 2016; Makovec, 2019; Nicholson, 2011; Taylor & Kuchel, 2009; von der Maase et al., 2005). However, clinical use of cisplatin is limited by cisplatin-induced nephrotoxicity (CIN), and there are limited alternate treatments for BCa (Nicholson, 2011; Oing et al., 2016; Sella & Kovel, 2012). Furthermore, despite the substantial increase in knowledge of the complex pathophysiology of CIN, there are currently no clinically approved therapeutic agents to prevent cisplatin-associated kidney injury (Baek et al., 2015; Kemp et al., 1996; Makimoto et al., 2018; Motwani, Kaur, & Kitchlu, 2022; Pabla & Dong, 2008; Shahbazi et al., 2015). This lack of treatment options is likely due to the fact that the various pharmacological, molecular, and genetic approaches identified as potential renoprotective strategies for CIN target pathways that are, at the same time, crucially involved in the cytotoxic effect of cisplatin on tumor cells (Miller et al., 2010; Pabla & Dong, 2008; Tang et al., 2023; Taylor & Kuchel, 2009). Thus, the advent of new therapeutic strategies to reduce CIN without affecting cisplatin’s anti-cancer effect is a critical unmet medical need.

Acute and repeated treatment of cisplatin result in drug accumulation in renal proximal tubular epithelial cells (Arany & Safirstein, 2003), and the severity of CIN in mice was recently correlated to the intratubular expression of the drug-metabolizing cytochrome P450 enzyme family member, CYP2E1 (Liu, Baliga, & Baliga, 2002; Liu & Baliga, 2003). In fact, the genetic deletion of CYP2E1 provided novel protection against CIN in *Cyp2e1^-/-^* mice (Liu & Baliga, 2003). This suggests that CYP2E1 activity could be a targetable primary event, which could reduce the occurrence of CIN. This warrants further investigation into the influence of CYP2E1 on CIN and the potential impact of its targeting on the treatment of BCa with cisplatin in male and female patients.

Evidence has indicated that male patients have a higher risk of developing CIN compared to females (Miyoshi et al., 2016; Mizuno et al., 2013). However, in other studies, female patients were shown to be more susceptible to CIN than males regardless of the type and stage of malignancy, dosage of cisplatin, drug-drug interactions, or the presence of comorbidities such as diabetes (Galfetti et al., 2020). Interestingly, data from stratified patients by age group or estrogen level (childbearing [< 45 years] and postmenopausal [> 55 years]) revealed that peri and postmenopausal women have a risk of developing CIN that is greater than men of the same age (Chen et al., 2017). This suggests that although the increased susceptibility of females to CIN may be multifactorial (Eshraghi-Jazi & Nematbakhsh, 2022), hormonal changes during menopause also play a role. In fact, Boddu et al. demonstrate that young female mice are significantly less susceptible to CIN than old female mice (Boddu et al., 2017), suggesting that estrogen production or low testosterone levels may have a protective effect on CIN. Currently, the mechanism underlying the sex-dependent susceptibility to CIN in patients is still poorly understood. CYP2E1 can be hormonally regulated (Konstandi, Cheng, & Gonzalez, 2013), and BCa mainly affects females when they are postmenopausal, with a median age of diagnosis of 73 years (Shariat et al., 2010; Taylor & Kuchel, 2009). So, the impact of CYP2E1 in premenopausal female mice’s susceptibility to CIN was evaluated in this study.

Our previous work demonstrated that 4-methylpyrazole (4MP; fomepizole), an FDA-approved antidote for methanol and ethylene glycol poisoning, also dose-dependently inhibits CYP2E1 activity (Akakpo et al., 2018). In fact, 50 μM of 4MP inhibited P450 enzyme activities in kidney homogenate by 72%, respectively (Akakpo et al., 2020). This suggests that 4MP-mediated inhibition of CYP2E1 could mitigate CIN. Yet, whether 4MP also prevents the toxicity of cisplatin in cancer cells is a critical gap in knowledge that needs to be addressed to confirm that targeting renal CYP2E1 with 4MP will not interfere with cisplatin anticancer effect via mechanisms that involve CYP2E1. Though not previously considered, the hepatoxic drug acetaminophen has recently been found to have anticancer potential (Bryan et al., 2023; Pingali et al., 2021). Additionally, our recent data suggests that 4MP does not impact the antitumor effect of a severe APAP overdose on commonly used 4T1 breast tumor and Lewis lung carcinoma tumor models (Bryan et al., 2024). This implies that 4MP is unlikely to impact cisplatin toxicity in BCa cells and, thus, could be a potential drug to use as a CYP2E1 inhibitor (Akakpo et al., 2024; Akakpo et al., 2019; Akakpo et al., 2018; Akakpo et al., 2023; Kang et al., 2020) to treat CIN in cancer patients. Hence, we hypothesize that CYP2E1 is central to the pathogenesis of CIN and that 4MP could be a promising drug candidate that can be rapidly repurposed to treat CIN and improve cisplatin tolerability and use in BCa patients. We measured renal CYP2E1 expression in male and female C57BL/6J mice, which we correlated to the development of CIN. To determine the potential relevance of CYP2E1 as a mechanistic target to treat CIN in cancer patients, we evaluated CYP2E1 expression in bladder tissue from male or female BCa patients (Stage I-III) and normal kidney tissue from male patients. Finally, we intervened with 4MP therapy targeted at CYP2E1 to determine if 4MP can prevent CIN in the mouse as well as in normal primary human kidney cells without interfering with cisplatin’s anti-cancer effects in human BCa cells.

## MATERIALS AND METHODS

### Experimental animals, acute kidney injury model

All animal procedures were approved by the Institutional Animal Care and Use Committee of the University of Kansas Medical Center. Experiments were performed following the guidelines of the National Research Council for the Care and Use of Laboratory Animals. In this study, 8 to 12-week-old male and female C57BL/6J mice were purchased from Jackson Laboratories (Bar Harbor, ME). Animals were acclimated to a temperature-controlled room with a 12-hour light/dark cycle, food, and water ad libitum for 3–5 days. For the acute kidney injury model, male and female mice were injected intraperitoneal (IP) with a single acute dose of 20 mg/kg cisplatin (Sigma-Aldrich, St. Louis, MO) or saline and were euthanized 3 days later (n= 6 animals per group). Due to their higher susceptibility to CIN, some male mice were also treated with a repeated dose of 9 mg/kg cisplatin once a week for 4 weeks before being euthanized. In addition, male mice received either a co-treatment of 20 mg/kg cisplatin (IP) + 50 mg/kg 4MP (IP) or 9 mg/kg cisplatin (IP) + 50 mg/kg 4MP (IP) (Sigma-Aldrich, St. Louis, MO) (n= 6 animals per group). Both cisplatin and 4MP were dissolved in saline. The animals were euthanized by cervical dislocation under isoflurane anesthesia. Blood was drawn from the caudal vena cava into heparinized syringes and was centrifuged at 20,000×g for 3 min at 4°C. Kidney tissue was also collected for analysis.

### Histopathological analyses

Collected kidney tissues were fixed in 4% formaldehyde overnight, embedded in paraffin blocks, and sectioned at 5 µm thickness. We deparaffinized sections and subjected them to hematoxylin and eosin (H&E) staining to assess the degree of morphological changes using a Nikon Eclipse Ti2 inverted microscope. Further examination of proximal tubular necrosis, proximal tubule degradation, and tubular casts served as an indication of morphological damage to the kidney after treatment with cisplatin or the combination of cisplatin and 4MP.

### Western blotting

Tissue samples were homogenized in ice-cold isolation buffer (pH 7.4) containing 220 mM mannitol, 70 mM sucrose, 2.5 mM HEPES, 10 mM EDTA, 1 mM EGTA, and 0.1% bovine serum albumin. This was followed by total protein measurements with the BCA assay from Pierce Scientific (Waltham, MA). Western blotting was then carried out with a rabbit anti-CYP2E1 antibody (Abcam, Cat. # ab28146). Anti-rabbit IgG horseradish peroxidase coupled secondary antibodies (Santa Cruz Biotechnology) (1:5000 dilution) and the ECL kit from Amersham (Piscataway, NJ) were then used for the detection of protein bands by chemiluminescence on the LI-COR Odyssey imaging system (LI-COR Biosciences).

### Immunostaining of human tissue samples

BCa and bladder normal tissue microarray, containing 35 urothelial carcinomas, 4 squamous cell carcinoma, 1 adenocarcinoma, 10 normal bladder tissue of male and female patients, were obtained from TissueArray (Array number BL1002b Derwood, MD). Normal human kidneys (NHK) from male cadavers, unsuitable for transplantation, were obtained from the Midwest Transplant Network (Kansas City, KS). Immunohistochemical staining of tissue samples was performed on paraffin-embedded sections cut at 5 μm thickness, deparaffinized and dehydrated. After sections were washed in phosphate buffer saline (PBS), they were blocked with 5% normal goat serum to reduce nonspecific reactions, followed by an overnight incubation step with the primary anti-CYP2E1 antibody (Cat. # ab28146, Abcam, Boston, MA). After 24 hours of incubation, sections were washed in PBS, followed by a 30-minute application of SignalStain Boost Detection Reagent (Cell Signaling Technology, Rabbit no. 8114), followed by SignalStain DAB Chromogen detection (Cat. # 8059, Cell Signaling Technology, Danvers, MA) as per manufacturer’s instructions. After counterstaining with hematoxylin, slides were imaged on the Nikon Eclipse Ti2 inverted microscope.

### Cell Culture

Male NHK cells were generated by the PKD Biomarkers and Biomaterials Core in the Kansas PKD Research and Translational Core Center at KUMC, as described previously (Wallace & Reif, 2019). Briefly, cells were seeded on collagen-coated plates in DMEM/F12 + P/S + ITS + 10% fetal bovine serum (FBS) and allowed to attach and proliferate in a humidified 5% CO2 incubator (at 37°C) for 2 days. The adherent NHK cells were washed with sterile PBS, and a starvation media consisting of DMEM/F12 + P/S + ITS + 0.5% FBS was added for 3 hours. NHK cells were treated with 20 µM cisplatin with or without 5 mM 4MP for 24 h. All drugs were dissolved in saline.

HTB9 BCa cells were acquired from the American Type Culture Collection (Manassas, VA). The cellular identity of HTB9 cells was confirmed twice yearly via STR analysis while in culture (University of Arizona Genetics Core). HTB9 cells were cultured as described previously (Woolbright et al., 2023). Briefly, cells were grown in DMEM supplemented with 10% FBS and 1% penicillin/streptomycin. Cells were incubated at 37 ° C with 5% C0_2_. At 24 h post-seeding, the media was changed until cells were approximately 90% confluent (Flaig et al., 2009). Cells were washed with sterile PBS and treated for 24 h with either cisplatin, 4MP, or both at the same concentrations used with kidney cells. All drugs were dissolved in saline.

### Biochemical measurements

Plasma samples were used to assess the extent of kidney injury by measuring blood urea nitrogen (BUN) and creatinine with QuantiChrom™ Assay kits from BioAssay Systems (Hayward, CA). NHK and HTB9 cell viability was assessed using an MTT (3- [4,5-dimethylthiazol-2-yl]-2,5 diphenyl tetrazolium bromide) cell proliferation and cytotoxicity detection assay (van Meerloo, Kaspers, & Cloos, 2011) following manufacturer instructions (Millipore Sigma; 11465007001). Cytochrome P450 enzyme activity was assessed in kidney homogenates using a fluorogenic substrate 7-ethoxy-4-trifluoromethyl coumarin (7-EFC) (Buters, Schiller, & Chou, 1993) (Invitrogen, Carlsbad, CA), as described previously (Akakpo et al., 2020).

### The Cancer Genome Atlas Program (TCGA) analysis

Data was acquired from the UALCAN website (http://ualcan.path.uab.edu). UALCAN is a comprehensive, user-friendly, and interactive web resource for analyzing cancer OMICS data (Chandrashekar et al., 2017; Chandrashekar et al., 2022). UALCAN allowed us to analyze the relative expression of CYP2E1 across tumor and normal samples, as well as in various tumor subgroups based on individual cancer stages, tumor grade, race, or other clinicopathologic features; graphs were processed using the UALCAN website.

### Statistical analysis

All statistical analyses were performed in SPSS Statistics 25 (IBM Co., Armonk, NY). One-way analysis of variance (ANOVA), followed by Student-Newman-Keul’s test, was used to test the significance between multiple groups. Statistical analysis between two groups was performed with the student’s two-tailed t-test. If the data were not normally distributed, the Kruskal-Wallis test (non-parametric ANOVA) followed by Dunn’s Multiple Comparison Test was used. Differences with values of p< 0.05 were considered statistically significant.

## RESULTS

### C57BL/6J male mice are more susceptible to cisplatin nephrotoxicity

To assess renal damage after cisplatin treatment, male and female C57BL/6J mice were treated with cisplatin (20 mg/kg i.p.) or saline (control group) for 3 days. Cisplatin induced a significant elevation of BUN and creatinine levels in male mice compared to control by day 3 of treatment (Figure 1A&B), but these changes were absent in the female mice. Cisplatin also induced damage in renal tissue of male mice treated with cisplatin, as indicated by varying degrees of tubular injury, including tubular dilation (black star) or rupture and tubular necrosis, as well as cast formation (black arrow), (x200) (Figure 1C). However, cisplatin-treated female kidney tissue shows normal renal corpuscle, glomerulus, and bowman’s capsule (yellow arrow) similarly to control male and female tissue. The histological evaluation of renal tissue was consistent with renal function markers. These observations demonstrate that C57BL/6J male mice are more susceptible to the nephrotoxic effects of cisplatin.

**Figure 1:**
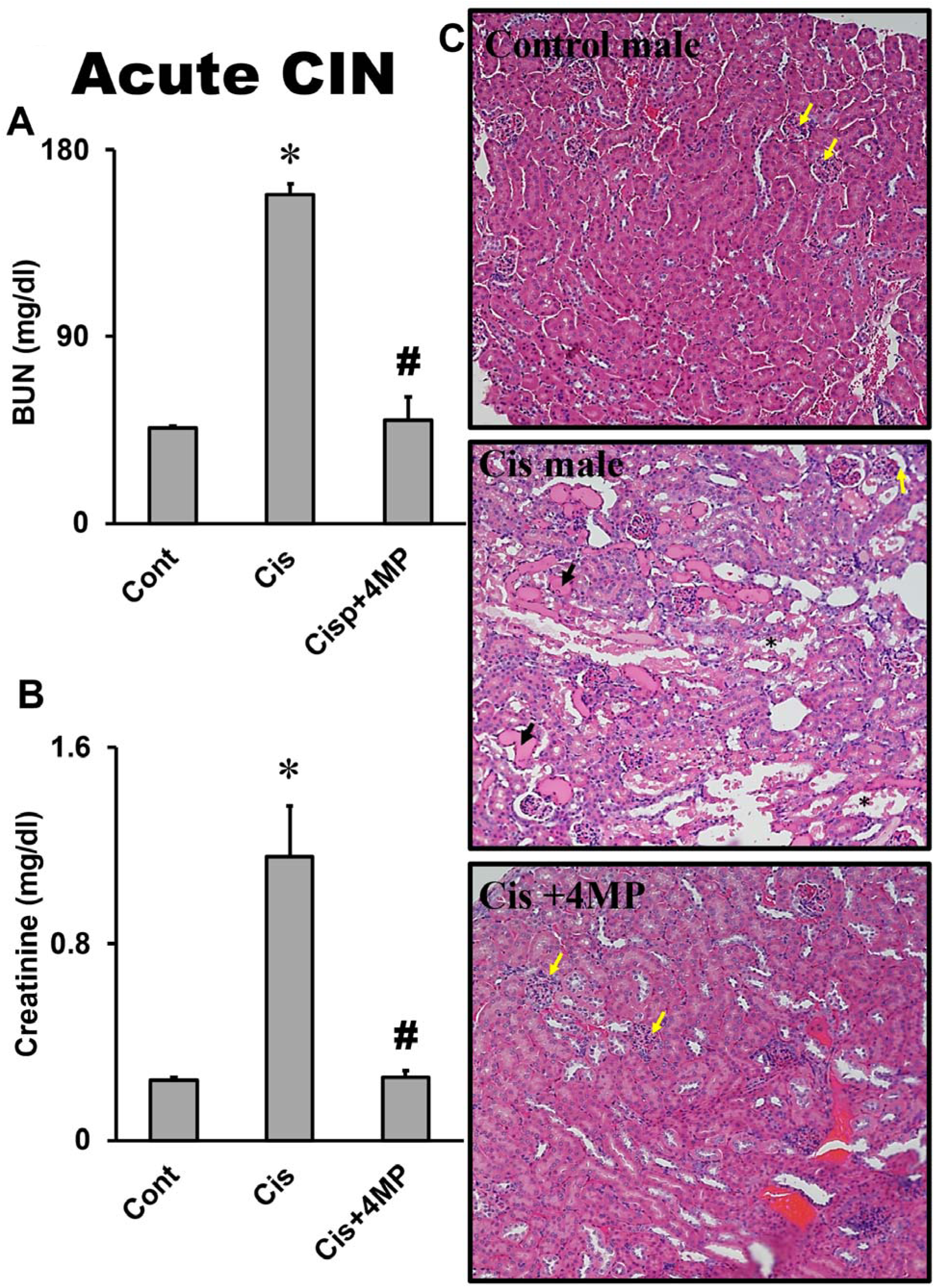
Sex-specific susceptibility to cisplatin-induced nephrotoxicity in C57BL/6J mice. C57BL/6J mice were treated with 20 mg/kg cisplatin or saline (control) for 3 days to collect blood and kidney tissue samples. Graphs are means ± SEM for plasma Blood Urea Nitrogen (BUN) (A) and creatinine levels (B). n= 6 animals per group. *p< 0.05 (compared to male control). Tissue sections were stained for H&E to visualize kidney morphology. Representative images of H&E staining of kidney sections (C) in control show normal renal corpuscle, glomerulus, and bowman’s capsule (yellow arrow). In contrast, tissue from cisplatin-dosed mice shows dilated tubules (black star) and cast formation (black arrowhead) (x200).

### Female mice have reduced renal CYP2E1 compared to male mice

CYP2E1 expression and activity have recently been demonstrated to be predominantly localized to the ER of human kidneys (Arzuk, Tokdemir, & Orhan, 2022). We and others have shown that renal proximal tubular epithelial cells of male kidneys predominantly express CYP2E1 in the ER (Akakpo et al., 2023 ; Liu & Baliga, 2003). Since male *Cyp2e1^-/-^*mice are resistant to CIN (Liu & Baliga, 2003), we examined whether CYP2E1 protein is expressed in the kidneys of female mice. Consistent with previous reports (Hoivik et al., 1995), we did not detect any expression of CYP2E1 in the kidneys of female mice when compared to male mice (Figure 2A). These data suggest that the lack of renal CYP2E1 expression may be the reason for the resistance of female mice to CIN. To test whether cisplatin could influence CYP2E1 activity, we assessed cytochrome P450 enzyme activity *in vitro* in a kidney homogenate from a male mouse using the fluorogenic substrate 7-EFC (Figure 2B). Cisplatin dose-dependently increased CYP2E1 activity, suggesting a direct effect of cisplatin on CYP2E1 activity in male mice but not female mice. We think that this difference contributes to the sex-specific differences in CIN.

**Figure 2:**
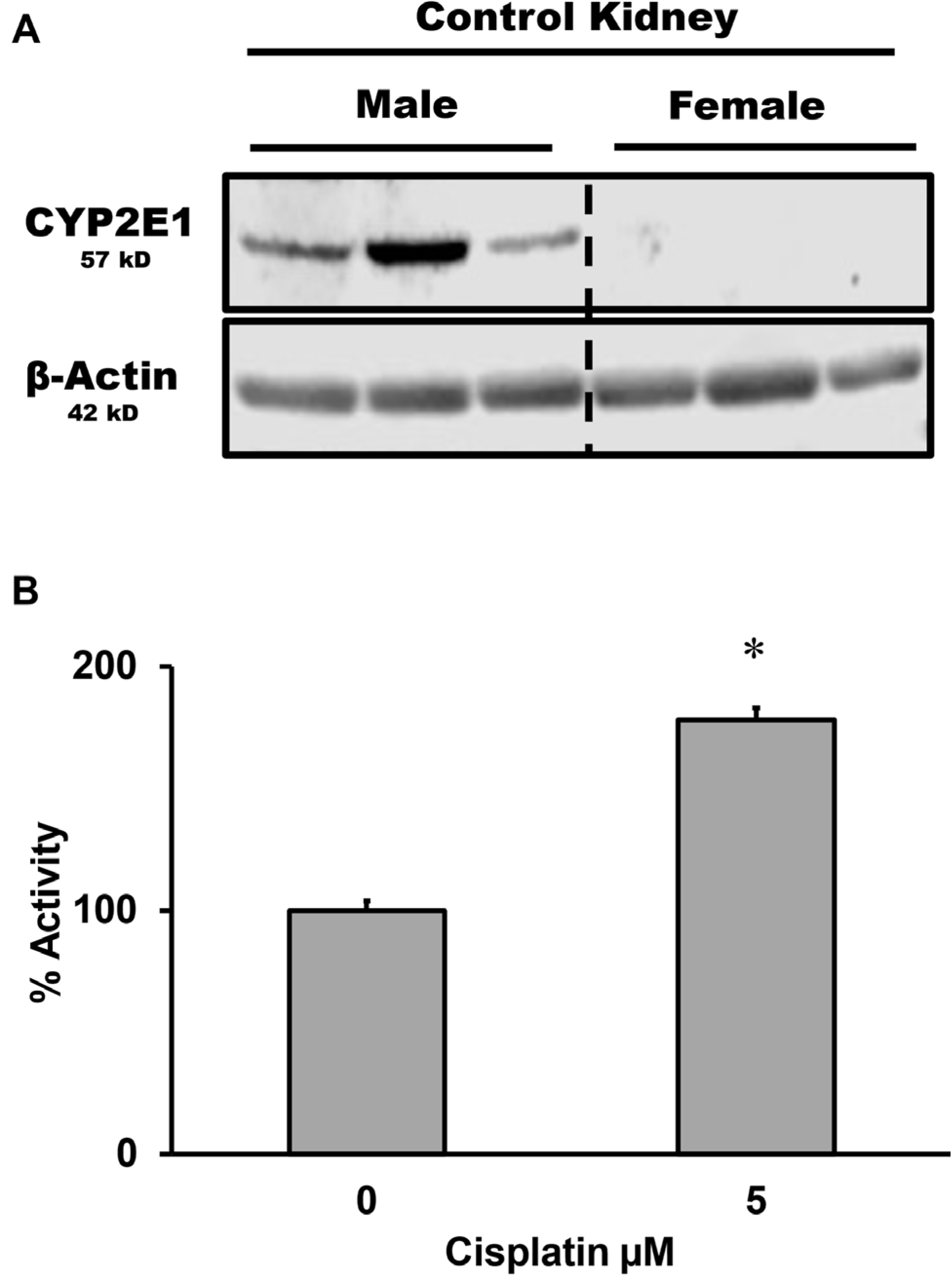
Expression of CYP2E1 is lower in the kidneys of female mice. **(A)** Kidney homogenates from untreated C57BL/6J male and female mice were subjected to western blot analysis to assess baseline expression of CYP2E1 (B) Cytochrome P450 enzyme activities were assayed in kidney homogenate using the fluorogenic substrate, 7-ethoxy-4-trifluoromethyl coumarin (7-EFC), which detects CYP2E1 and CYP1A2 activities, in the presence or absence of 5 μM 4MP. The enzyme activity of 178.08 RFU/mg protein/min was normalized to 100% of control activity. Data represent means ± SEM of 3 separate measurements. *P< 0.05 (compared to the respective control).

### 4MP prevented the kidney injury induced by cisplatin

To determine if targeting CYP2E1 may prevent CIN, male mice were treated with the FDA-approved drug 4MP, which inhibits CYP2E1 (Figure 2). Male C57BL/6J mice were treated with a single acute dose of 20 mg/kg cisplatin and either 50 mg/kg 4MP or saline (control). Then, kidney injury was evaluated after 3 days to determine the impact of CYP2E1 inhibition on acute cisplatin exposure. Additional mice were treated with the repeated dosing regimen of 9 mg/kg cisplatin and 50 mg/kg of 4MP or saline once per week for 4 weeks. Blood and kidney tissue samples were then collected to assess the extent of kidney injury. Our data show that after both the acute and repeated cisplatin treatments, male mice developed severe kidney injury as indicated by elevated plasma BUN (Figures 3 and 4A) and creatinine levels (Figures 3 and 4B) when compared to saline-treated control. This is consistent with the histological analysis performed on tissue samples collected after the acute and repeated cisplatin treatment, which shows extensive tubular dilation, tubular necrosis (black star), and cast formation (black arrows) (Figures 3 and 4C). Interestingly, the co-treatment of cisplatin with 4MP almost completely prevented cisplatin toxicity during cisplatin treatment, as indicated by the lower plasma BUN and creatinine levels, as well as the absence of detectable tubular dilation and necrosis or cast formation (Figures 3 and 4). These data suggest that the inhibition of CYP2E1 with 4MP blocked the development of CIN.

**Figure 3:**
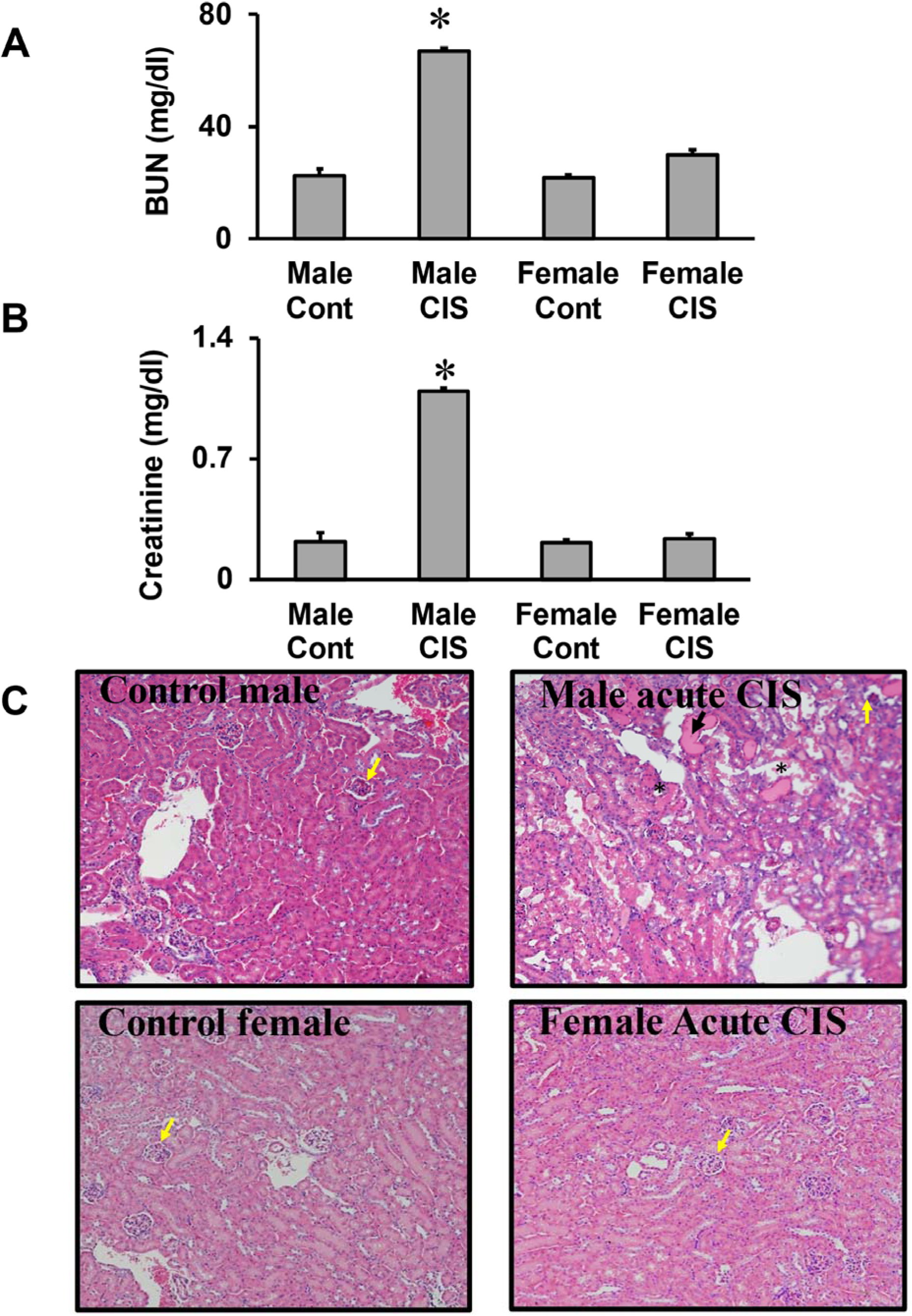
4MP protects against cisplatin-induced acute kidney injury in C57BL/6J mice. C57BL/6J mice were treated with 20 mg/kg cisplatin and either 50 mg/kg 4MP or saline (control) for 3 days to collect blood and kidney tissue samples. Graphs are means ± SEM for plasma blood urea nitrogen (BUN) (A) and creatinine levels (B). n= 6 animals per group. *p< 0.05 (compared to female control). #p<0.05 (compared to cisplatin). Tissue sections were stained for H&E to visualize kidney morphology. Representative images of H&E staining of kidney sections (C) in control and cisplatin + 4MP treatment show normal renal corpuscle, glomerulus, and Bowman’s capsule (yellow arrow). In contrast, tissue from cisplatin-dosed mice shows dilated tubules, tubular necrosis (black star), and cast formation (black arrowhead) (x200).

**Figure 4:**
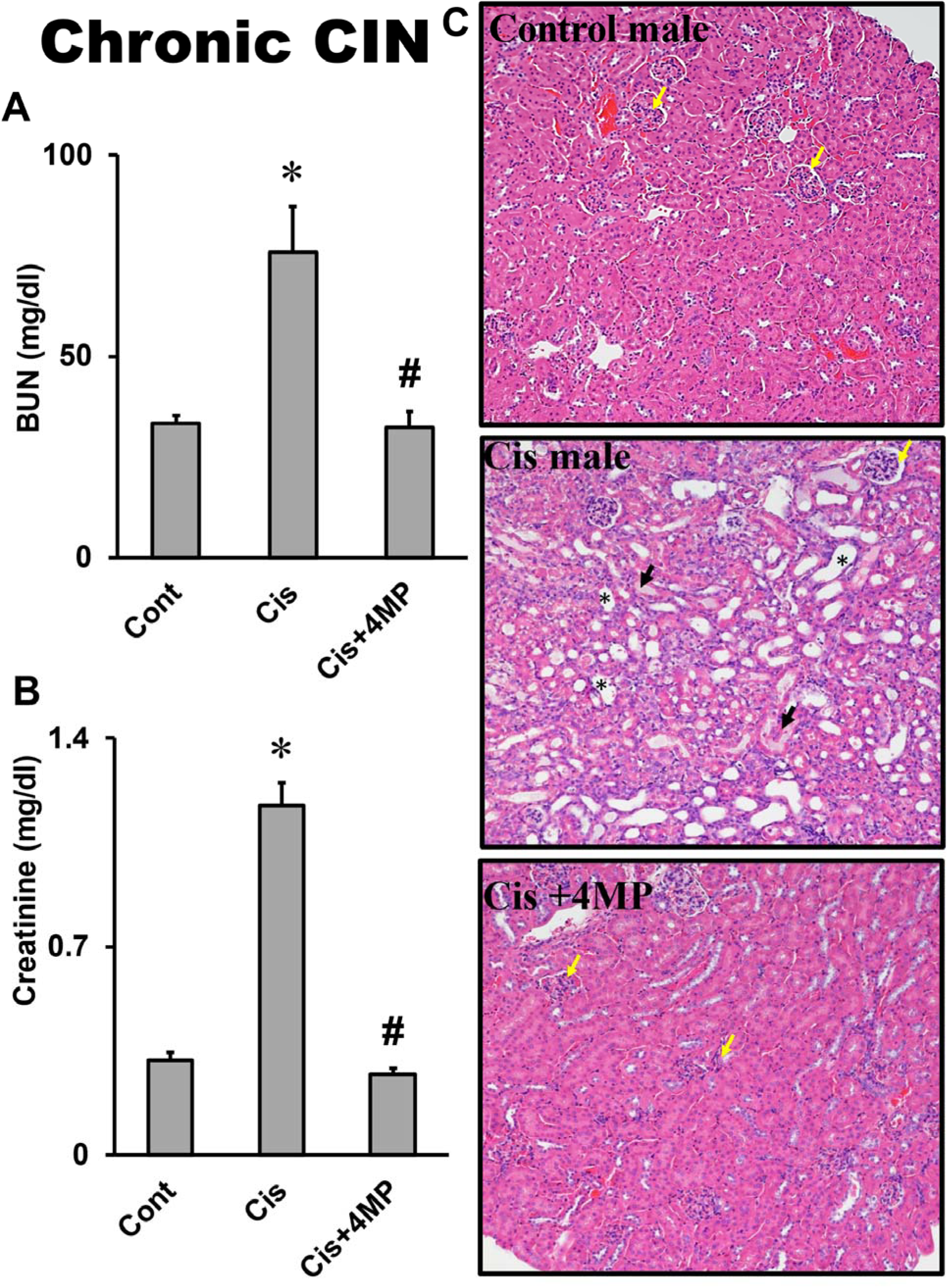
4MP protects against cisplatin-induced chronic kidney injury in C57BL/6J mice. C57BL/6J mice were treated with 9 mg/kg cisplatin and 50 mg/kg of saline control once per week for 4 weeks to collect blood and kidney tissue samples. Graphs are means ± SEM for plasma blood urea nitrogen (BUN) (A) and creatinine levels (B). n= 6 animals per group. *p< 0.05 (compared to female control). #p<0.05 (compared to cisplatin). Representative images of H&E staining of kidney sections (C) in control and cisplatin + 4MP treatment show normal renal corpuscle, glomerulus, and bowman’s capsule (yellow arrow). In contrast, tissue from cisplatin-dosed mice shows dilated tubules, tubular necrosis (black star), and cast formation (black arrowhead) (x200).

### CYP2E1 expression and inhibition in patients with BCa

Prior to the initiation of chemotherapy, BCa patients must have acceptable creatinine levels or calculated creatinine clearance, indicating the absence of kidney injury (Nicholson, 2011; Taylor & Kuchel, 2009). Thus, we evaluated CYP2E1 expression levels in BCa patients using the extensive TCGA database via UALCAN (Chandrashekar et al., 2017; Chandrashekar et al., 2022) (Figures 5 and 6). Our results indicate a trend toward the downregulation of CYP2E1 gene expression in patients with BCa compared to normal control tissue (Figure 5A) at all BCa cancer stages (Figure 5B), however, this finding does not reach statistical significance. Interestingly, this trend toward the downregulation of CYP2E1 gene expression was independent of race (Figure 6A) or sex (Figure 6B). While the UALCAN data suggest a potential downregulation of CYP2E1 gene expression in BCa patients, further analysis with larger patient sample sizes is warranted to confirm this observation. To verify whether the trend observed with CYP2E1 gene expression correlates with protein expression in BCa tissue, we subsequently evaluated CYP2E1 expression levels in BCa and normal bladder tissue microarrays containing 35 urothelial carcinomas, 4 squamous cell carcinomas, 1 adenocarcinoma and 10 normal bladder tissues from males and females that we compared to the normal kidney from male patients.

**Figure 5:**
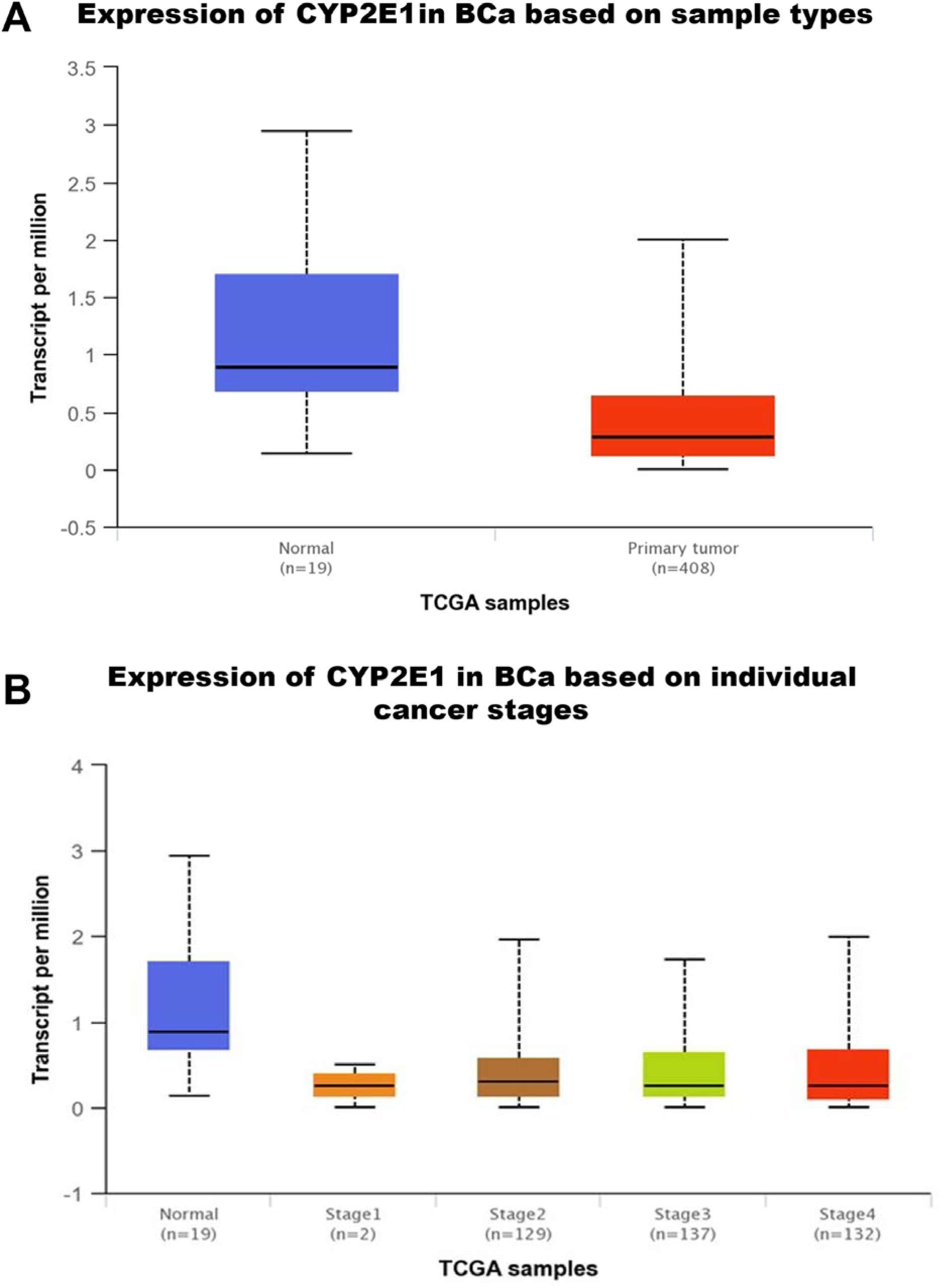
CYP2E1 expression in patients with primary BCa tumor. The gene expression and clinical patient data were downloaded from TCGA and processed to generate graphical outputs. Samples were categorized using clinical patient data, and a boxplot was generated for the expression level of CYP2E1 (A). Samples were divided into individual cancer stage I, stage II, stage III, and stage IV groups, and a boxplot was generated for the expression level of CYP2E1 (B).

**Figure 6:**
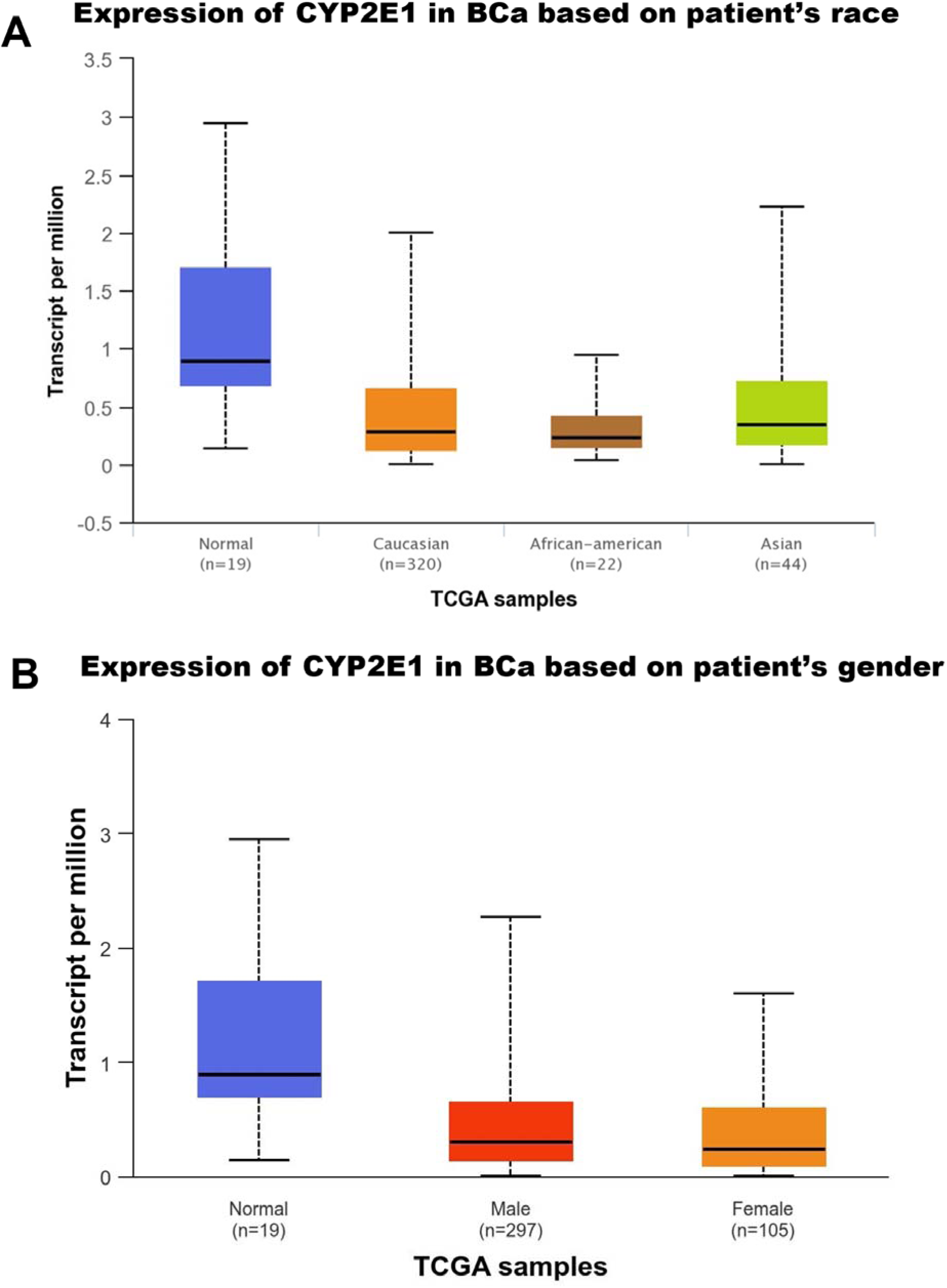
Race and sex and differences in CYP2E1 expression of BCa human patients. The gene expression and clinical patient data were downloaded from TCGA and processed to generate graphical outputs. Samples were divided using patient race information into Caucasian, African American, and Asian groups, and a boxplot was generated for the expression level of CYP2E1 (A). Samples from male and female patients were grouped separately to account for gender differences, and a boxplot was generated to evaluate the expression level of CYP2E1 (B).

Immunohistochemistry staining for CYP2E1 revealed appreciable signals in kidney tissue (Figure 7A), consistent with our previous report (Akakpo et al., 2023). However, CYP2E1 expression was undetectable in normal bladder tissue or BCa samples at all stages (Figure 7 B & Supplementary Figure 1). This indicates that targeting CYP2E1 with 4MP is unlikely to interfere with the antineoplastic effect of cisplatin in BCa patients while protecting kidneys against CIN.

**Figure 7:**
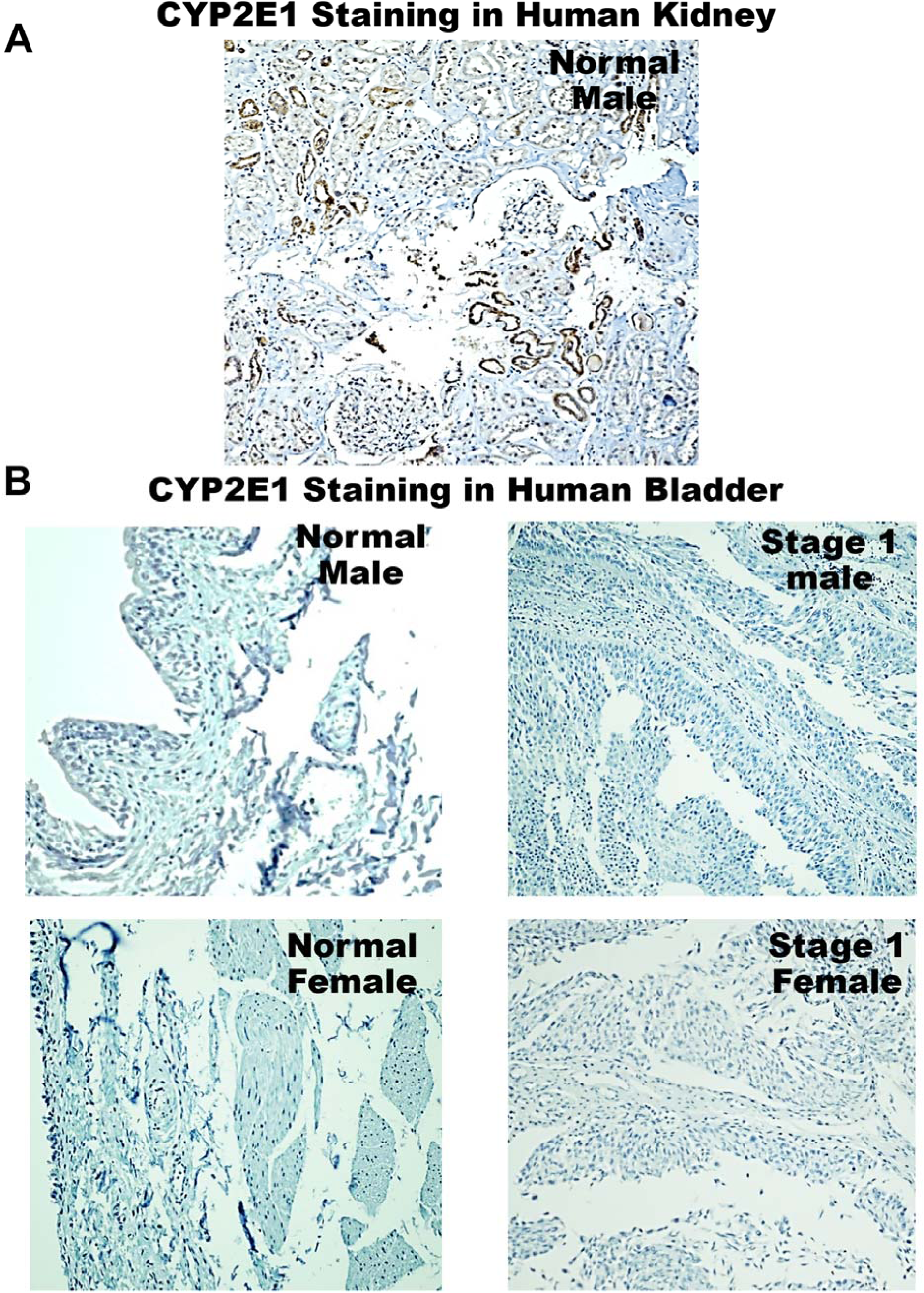
CYP2E1 expression in the kidney and bladder of human tissue sections. The CYP2E1 expression level was evaluated by immunohistochemistry staining of human BCa and normal bladder tissues as well as kidney tissue sections. Bladder images are representative of male and bladder carcinoma or normal tissue derived from a tissue microarray, containing 35 urothelial carcinoma, 4 squamous cell carcinoma, 1 adenocarcinoma, 10 normal bladder human tissue (A). The kidney tissue section image is representative of 3 male donors per group. (magnification: x200) (B).

### CYP2E1 inhibition protects renal epithelial cells, but not BCa cells, from cisplatin-induced cell death

To confirm the above assumption derived from analysis of the TCGA data and immunostaining bladder tissue and further evaluate whether targeting CYP2E1-mediated mechanisms of cisplatin-induced renal proximal tubular cell death with 4MP are relevant to humans, we treated NHK cells or HTB9 BCa cells with 5, 10, or 20 µM cisplatin with or without 5 mM 4MP for 24 h (figure 8). The data shows that cisplatin induced a dose-dependent cell death in these human kidney cells, which was significantly reduced with 4MP co-treatment (Figure 8A). Interestingly, the presence of 4MP did not affect cisplatin-induced HTB9 BCa cell death, as indicated by a similar loss of cell viability in BCa cells exposed to cisplatin alone or cells exposed to the combination of cisplatin + 4MP (Figure 8B). This data demonstrates that 4MP-mediated CYP2E1 inhibition appears to be an attractive renoprotective strategy to prevent cisplatin nephrotoxicity without affecting the chemotherapeutic effects of cisplatin on BCa.

**Figure 8:**
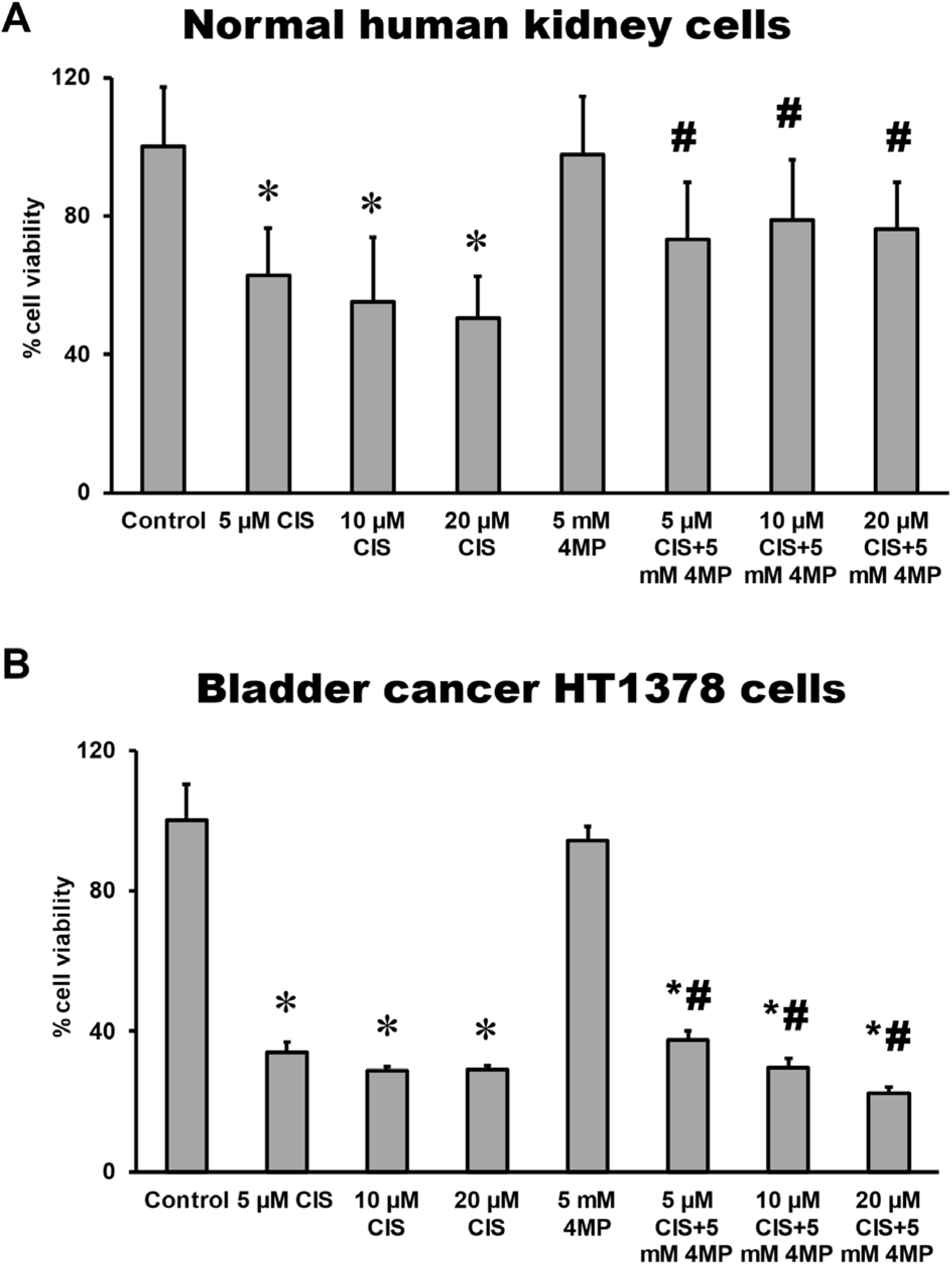
Effect of 4MP in cisplatin-treated NHK cells and bladder cancer HTB9 cells. Cells were treated with 20 µM cisplatin alone or in combination with 5 mM 4MP. % MTT cytotoxicity assay was performed to determine % cell viability (A). Bars represent means ± SEM for three separate experiments. *p < 0.05 versus control; #p < 0.05 versus cisplatin alone treatment. (CIS=cisplatin).

## DISCUSSION

The purpose of this study was to investigate whether targeting CYP2E1 reduces CIN without interfering with its antineoplastic effects in BCa. The efficacy of the FDA-approved drug 4MP against CIN was tested in normal human kidney cells and human BCa cells as well as *in vivo* in a murine model.

### Bioactivation of cisplatin in the development of CIN

The pathophysiology of cisplatin-induced acute kidney injury (AKI) involves various molecular and cellular mechanisms, including increased oxidative stress (Liu & Baliga, 2003; Yamamoto et al., 2024; Zhang et al., 2020), inflammation (Faubel et al., 2007; Lu et al., 2008; Ramesh & Reeves, 2002; Zhang et al., 2008), and apoptosis (Lee et al., 2001; Lieberthal, Triaca, & Levine, 1996; Wei et al., 2007). However, the mechanistic events that trigger CIN are not well-established. The renal metabolism and bioactivation of cisplatin to a nephrotoxin have been suggested to play a major role in the pathophysiology of CIN (Townsend, Deng, et al., 2003). After accumulation in the proximal tubule, cisplatin can bind to intrarenal GSH in a reaction mediated by glutathione-S-transferase to form glutathione–cisplatin conjugates (Townsend, Deng, et al., 2003; Townsend & Hanigan, 2002; Townsend, Marto, et al., 2003; van Bladeren, 2000). Extracellular transport of the GSH conjugate and subsequent consecutive reactions catalyzed by γ–glutamyltranspeptidase (GGT) and aminodipeptidase (DP) result in the formation of cisplatin-cysteinyl-glycine and cisplatin-cys conjugates respectively (Hanigan et al., 1994; Hanigan et al., 2001; Townsend & Hanigan, 2002). The pyridoxal 5’-phosphate-containing enzyme, cysteine-s conjugate beta-lyase (Katayama et al., 2011; Zhang & Hanigan, 2003), then catalyzes the beta-elimination reactions with cisplatin-cys to generate a highly reactive electrophilic nephrotoxin (thiol-cisplatin) (Katayama et al., 2011; Sawers et al., 2014). The nephrotoxin binds to cellular macromolecules to ultimately trigger apoptotic and necrotic renal cell death (Lieberthal, Triaca, & Levine, 1996; Miller et al., 2010). Interestingly, the thiol-cisplatin is formed after the extracellular transport and metabolism of a cisplatin GSH conjugate by the cell surface enzymes, GGT, and DP (Hanigan et al., 1994; Townsend, Deng, et al., 2003; Townsend & Hanigan, 2002) (Figure 9). However, CYP2E1 is likely also toxicologically important to the pathology of CIN because its genetic deletion provides novel protection against CIN (Liu & Baliga, 2003). Recent results from our group indicate that CYP2E1 is expressed in the ER of renal proximal tubular cells of C57BL/6J male mice (Akakpo et al., 2023). This aligns with previous immunoelectron microscopy experiments with gold particles in mice (Liu & Baliga, 2003). Our in vitro data now shows that cisplatin increases the activity of CYP2E1 (Figure 2B), suggesting a potential direct interaction between the drug and the enzyme. Yet, whether this is replicated *in vivo* is unclear and requires further investigation. Importantly, extracellular transport of cisplatin-GSH is necessary to form cisplatin thiols, while CYP2E1 is expressed intracellularly (Akakpo et al., 2023; Liu & Baliga, 2003). Thus, the interaction of cisplatin with CYP2E1 would need to occur prior to the extracellular transport of cisplatin-GSH and subsequent metabolism with GGT and DP. This highlights the fact that the exact sequence of cisplatin-CYP2E1 interaction and its activation to cisplatin-thiol are still vague, and the primary event driving CIN remains to be determined.

**Figure 9:**
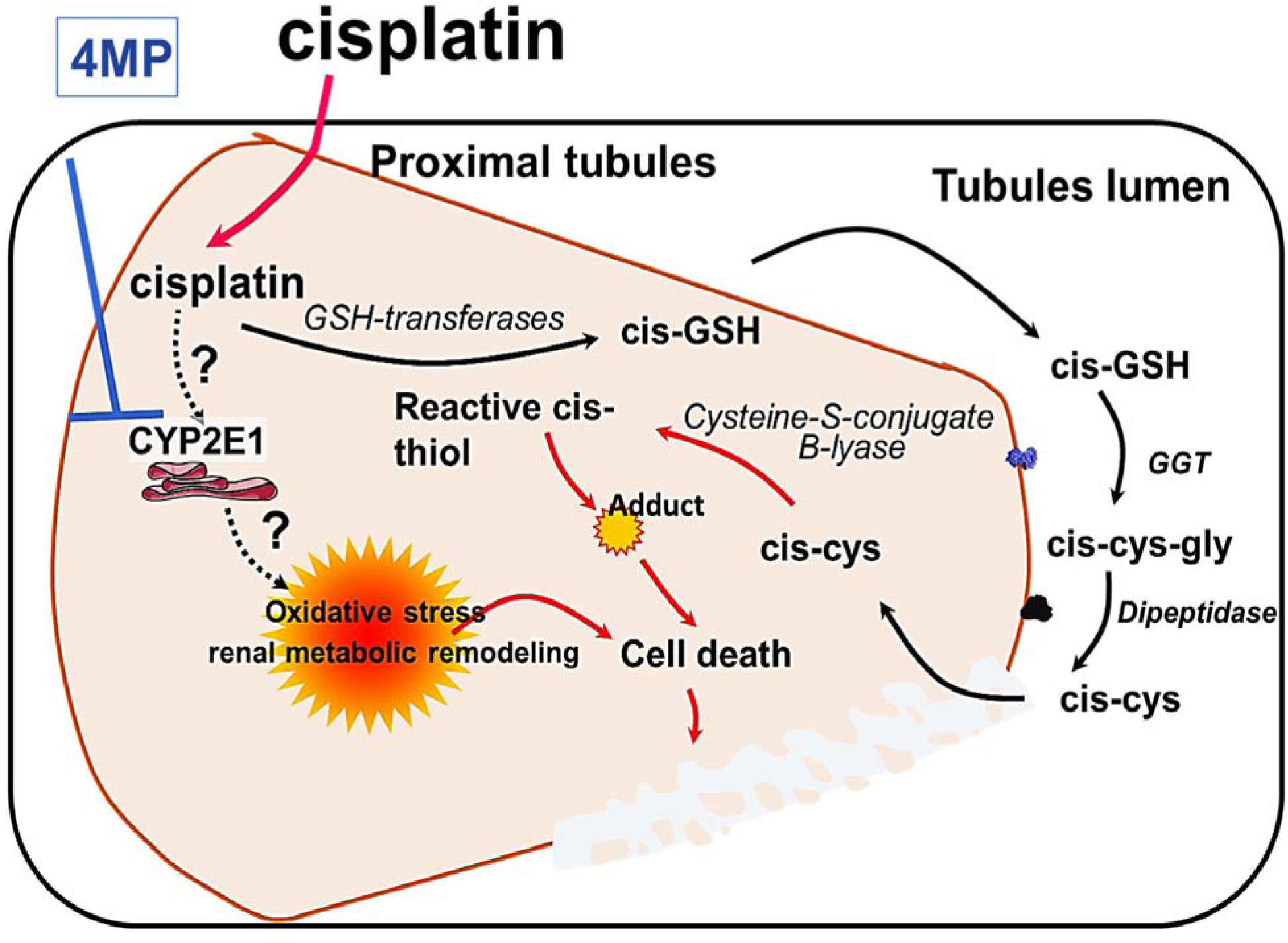
Effect of 4MP on the metabolism of cisplatin in the kidney. Cisplatin enters cells and binds to DNA, which is toxic to dividing BCa cells (A) but not to quiescent proximal tubular epithelial cells (PTECs) in the kidney (B). In PTECs, cisplatin forms cisplatin-glutathione conjugates (Cis-GSH), which is then cleaved to a cisplatin-cysteinyl-glycine conjugate (cis-cys-gly) by GGT expressed on the cell surface. Cis-cys-gly is further cleaved to a cisplatin-cysteine conjugate (cis-gly) by aminodipeptidase, also expressed on the cell surface. The platinum-cysteine conjugate is taken up into the cell, where it is converted to a highly reactive thiol by cysteine S-conjugate β-lyase (cis-thiol). The binding of the reactive thiol to essential proteins within the cell is toxic. Cisplatin also influences tubular cell death via unknown mechanisms that involve CYP2E1 expressed in the ER of the cell. 4MP protects PTECS against cisplatin-induced nephrotoxicity.

### CYP2E1’s influence on the sex-specific toxic renal effect of cisplatin

Our data indicates that male mice, in contrast to females, develop severe kidney injury after cisplatin treatment. This is consistent with a previous study demonstrating that young C57BL/6J female mice are also less susceptible to CIN compared to male mice (Boddu et al., 2017). In fact, most of the experimental studies have reported that female rodents are more resistant than males to both the acute and repeated cisplatin treatment (Eshraghi-Jazi & Nematbakhsh, 2022; Nematbakhsh et al., 2017; Shi et al., 2018). Thus, in animal models, sex differences have a significant influence on the severity of nephrotoxicity induced by cisplatin. This is clinically relevant because prior studies have highlighted sex-specific differences in the susceptibility to CIN in humans (Galfetti et al., 2020; Miyoshi et al., 2016; Mizuno et al., 2013). Notably, age-stratified patient data indicated that peri- and post-menopausal women have a higher risk of developing CIN compared to men of the same age (Chen et al., 2017). This aligns with animal models showing that aged female mice are more susceptible to CIN than young female mice (Boddu et al., 2017). Moreover, clinical epidemiology shows that BCa mainly affects post-menopausal women with a median age of diagnosis of 73 years (Shariat et al., 2010; Taylor & Kuchel, 2009). However, the mechanism underlying the sex-dependent susceptibility to CIN in patients is still poorly understood. Understanding this mechanism is crucial for optimizing treatment strategies and minimizing adverse effects. In this study, we provide evidence for a significant role of CYP2E1 in mediating the sex-specific differences in the susceptibility to CIN since female C57BL/6J mice lack renal CYP2E1 expression and are resistant to CIN. The data suggest that renal CYP2E1 expression in male mice may lead to enhanced bioactivation of cisplatin, resulting in a greater cisplatin nephrotoxic effect, which is not observed in female mice. This is also relevant to humans since CYP2E1 activity has been shown to be lower in female kidneys compared to males (Arzuk et al., 2018). Yet, our data do not rule out the possibility that additional confounding factors may contribute to the resistance of female mice to CIN. For instance, there are sex-related differences in the organic cation transporter (OCT2) (Urakami et al., 1999), which is responsible for cisplatin uptake into renal tubular cells and has significantly higher expression in male kidneys (Urakami et al., 1999), thereby increasing the accumulation and exposure to cisplatin in proximal tubules in males. Sex-specific differences have also been noted in the expression of GGT (Darnerud et al., 1991, Tanaka et al., 1992), with higher expression in males than in female mice, and this may facilitate the formation of cisplatin-thiol for a potential potentiation of CIN. Females also have lower oxidative stress (Kander, Cui, & Liu, 2017; Townsend et al., 2009) in response to cisplatin than males, and this variation may additionally play a role in the female mice’s resistance to CIN. This further supports an implication of CYP2E1 in CIN since the enzyme plays a significant role in oxidative stress (Lu et al., 2012; Qi et al., 2013), which is a well-documented contributor to CIN (Liu & Baliga, 2003; Yamamoto et al., 2024; Zhang et al., 2020). While the enzyme is constitutively expressed intracellularly in proximal tubules, it can also be upregulated in certain pathophysiologic conditions such as diabetes (Lucas et al., 1998), starvation (Mandl et al., 1995), and obesity (McCarver et al., 1998). Moreover, the sex-specific difference in susceptibility in humans could also be due to GSH metabolism (Wang, Ahn, & Asmis, 2020), but the estrous cycle regulation of CYP2E1 may also play a role (Konstandi, Cheng, & Gonzalez, 2013). Thus, the specific mechanism of the contribution of CYP2E1 to the resistance of female mice to CIN remains to be explored in further animal experiments.

### 4MP and the prevention of CIN clinically

The preceding discussion builds a case for CYP2E1 expression and activity being responsible for the higher susceptibility of kidneys to CIN. Furthermore, our earlier findings indicate that 4MP does not affect the antitumor efficacy of a severe acetaminophen overdose in commonly used tumor models, specifically the 4T1 breast tumor and Lewis lung carcinoma models (Bryan et al., 2024). Additionally, the absence of detectable CYP2E1 expression in human BCa tissue samples suggests that the mechanism by which cisplatin induces cell death in BCa cells is independent of CYP2E1. This contrasts with its role in kidney tissues and implies that therapeutic strategies targeting CYP2E1 may be beneficial in reducing CIN, the major use-limiting factor, in cancer patients without affecting the anti-cancer effect of cisplatin. BCa patients need to have acceptable kidney function prior to the initiation of cisplatin therapy (Nicholson, 2011).

Interestingly, our staining experiments show that BCa patients do not express CYP2E1 in tumors (Figure 7). Thus, targeting CYP2E1 appears to be a logical nephroprotective strategy to reduce the occurrence of CIN without interference with the antineoplastic effect of cisplatin. We provide evidence *in vivo* that inhibition of CYP2E1 activity by 4MP co-treatment with the acute or repeated cisplatin treatment protects male mice against CIN. To evaluate if the reduction of cisplatin nephrotoxicity by 4MP was specific to the kidney, we looked for potential reductions in the chemotherapeutic efficacy of cisplatin in human HTB9 BCa cells. Our experiments further revealed that while 4MP dose-dependently prevents CIN in NHK cells, it does not interfere with cisplatin toxicity in human BCa cells (Figure 8). Our data suggests that 4MP, an already FDA-approved drug in clinical use, is a promising drug candidate that could be rapidly repurposed to prevent CIN without interfering with cisplatin’s antineoplastic effect. However, further translational investigations in clinically relevant BCa cancer mouse models are required to provide strong evidence for the potential benefit of 4MP in the clinic.

In conclusion, cisplatin is the most active chemotherapeutic used in the treatment of various cancers (Makovec, 2019) and is considered first-line therapy in BCa. However, nephrotoxicity, characterized by extensive tubular injury and rapid loss of renal function, occurs in around 30% of patients who are receiving cisplatin chemotherapy. Unfortunately, there is currently no FDA-approved treatment to reduce or prevent CIN, and there are limited 2nd line treatments for BCa. To the best of our knowledge, this is the first study to demonstrate the potential protective effects of 4MP against the likely primary mechanism of CIN, where the influence of cisplatin on CYP2E1 activity drives CIN. It would suggest that *in situ* activation of cisplatin within the kidney is a primary event that precedes cisplatin-thiol activation in the etiology of CIN. Early treatment with 4MP prior to initiation of cisplatin could be a viable prophylactic therapeutic option to prevent CIN clinically because it could significantly prevent kidney injury after the acute or repeated cisplatin treatment dosing regimen by inhibiting CYP2E1 without interfering with cisplatin’s toxic effects on cancer cells. Future research will focus on clarifying the precise molecular mechanisms by which CYP2E1 contributes to sex-specific differences in cisplatin nephrotoxicity. Future work will also determine whether 4MP can reverse injury progression after the acute or repeated treatments of cisplatin. This will help create a rationale for the clinical use of 4MP as a feasible treatment to efficiently promote the antineoplastic effect of cisplatin in bladder cancer cells while preventing its toxic side effects on healthy kidney cells.

## ACKNOWLEDGEMENTS

Funding for this study was provided by the Leo and Anne Albert Charitable Trust (JAT). This work was partly funded by a postdoctoral fellowship (J.Y.A.) from the CTSA grant funded by NCATS at the University of Kansas for Frontiers: University of Kansas Clinical and Translational Science Institute No. TL1TR002368. Primary cultures of human kidney cells were provided by the PKD Biomarkers and Biomaterials Core Laboratory in Kansas PKD Research and Translational Core Center (RTCC; NIH U54 DK126126) at KUMC and the national PKD Research Resource Consortium (PKD-RRC).

## FOOTNOTES

**The authors declare no conflict of interest.**

