## Supplemental figure 1 for "4-Methylpyrazole-mediated inhibition of Cytochrome P450 2E1 protects renal epithelial cells, but not bladder cancer cells, from cisplatin toxicity"

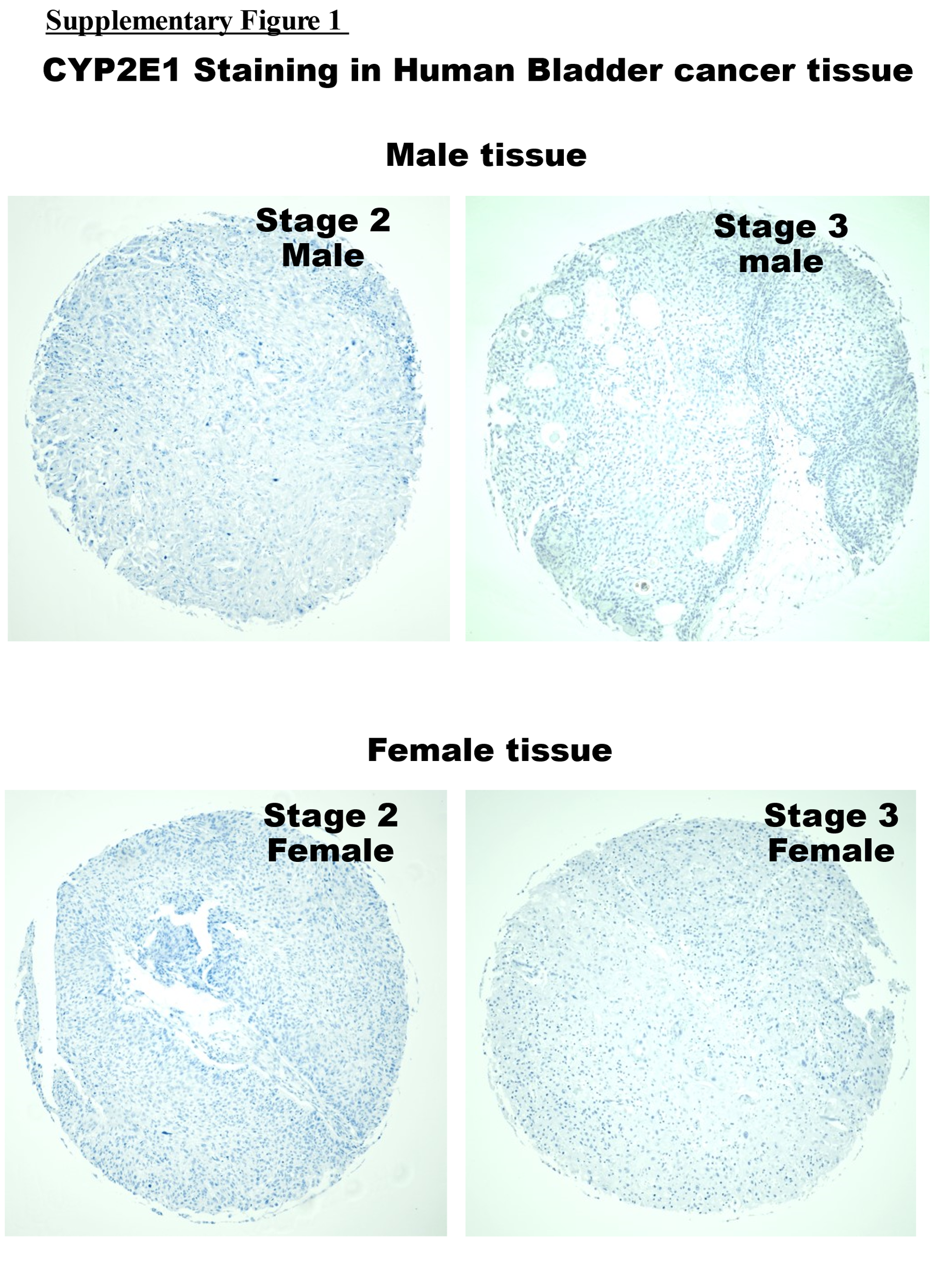

**Supplementary Figure 1: CYP2E1 expression in stage 2 and 3 bladder cancer tissue sections.** CYP2E1 in the kidney (A) (x200). Immunohistochemistry staining of Cyp2E1 male kidney tissue (A). Immunohistochemistry staining of Cyp2E1 in normal and stage 1 bladder tissue sections (B) (magnification: x200. (Kidney image Is representative of 3 male donors per group. Bladder images are representative of male and bladder carcinoma and normal tissue microarray, containing 35 urothelial carcinoma, 4 squamous cell carcinoma, 1 adenocarcinoma, 10 normal bladder human tissue obtained. Unstained microarray slides were obtained from TissueArray (BL1002b) and stained for CYP2E1)
